# Rbfox1 mediates cell-type-specific splicing in cortical interneurons

**DOI:** 10.1101/305904

**Authors:** Xavier Hubert Jaglin, Brie Wamsley, Emilia Favuzzi, Giulia Quattracolo, Maximiliano José Nigro, Nusrath Yusef, Alireza Khodadadi-Jamayran, Bernardo Rudy, Gordon Fishell

**Author notes:** These authors contributed equally (Xavier Hubert Jaglin, Brie Wamsley).

## Abstract

Cortical interneurons display a remarkable diversity in their morphology, physiological properties and connectivity. Elucidating the molecular determinants underlying this heterogeneity is essential for understanding interneuron development and function. We discovered that alternative splicing differentially regulates the integration of somatostatin- and parvalbumin-expressing interneurons into nascent cortical circuits through the cell-type specific tailoring of mRNAs. Specifically, we identified a role for the activity-dependent splicing regulator Rbfox1 in the development of cortical interneuron subtype specific efferent connectivity. Our work demonstrates that Rbfox1 mediates largely non-overlapping alternative splicing programs within two distinct but related classes of interneurons.

## Introduction

Cellular diversity within the central nervous system has evolved to support intricate neuronal circuits and a broad range of complex behaviors. Cortical interneurons (cINs) are emblematic of this diversity. While representing only a small proportion of the cells within the cortex, discrete cIN subclasses are critical for supporting network functions. One remarkable aspect of this diversity is their subtype specific synaptic connectivity (Blackstad and Flood, 1963). While the Parvalbumin (PV)+ cINs innervate the perisomatic region of pyramidal excitatory neurons, Somatostatin (SST)+ cINs preferentially target their distal dendrites. In addition to being useful features for cell classification, these differences in subcellular targeting are critical to their function (Tremblay et al., 2016). However, whether these distinct features are genetically hard-wired, imposed by environmentally-driven mechanisms (such as early neuronal activity) or some combination of the two is poorly understood (Wamsley and Fishell, 2017).

While there is no doubt that transcription regulation is involved in cIN specification and maturation (reviewed in Batista-Brito and Fishell, 2009) it alone seems unlikely to account for the full range of functional diversity within cIN subtypes. Consistent with this idea, neurons utilize a variety of other modes of gene expression regulation to enhance their molecular diversity. These include the differential usage of transcription start sites, alternative RNA splicing, polyadenylation and editing (Maniatis and Tasic, 2002). Indeed, about 90-95% of transcripts from human genes undergo alternative splicing (AS) (Johnson et al., 2009; Pan et al., 2008; Wang et al., 2008a). Recent genome-wide investigations of AS in mice have highlighted its prevalence during development within various regions of the nervous system (Dillman et al., 2013; Pan et al., 2008; Wang et al., 2008b; Yan et al., 2015). Specifically, AS participates in cell fate decisions during cortical neurogenesis (Linares et al., 2015; Zhang et al., 2016), as well as synaptogenesis and synaptic plasticity (reviewed in Raj and Blencowe, 2015; Vuong et al., 2016).

In the present study, we find that AS events play a central role within SST+ and PV+ cINs during their integration into the cortex. Specifically, we discovered that the number of AS events within these cell types varies as they mature. Moreover, our work indicates that Rbfox1 function is essential in regulating distinct AS events within developing PV+ and SST+ cINs. Together our findings demonstrate that Rbfox1-mediated AS is essential for the genetic regulation of developing cIN connectivity.

## Results

### Alternative splicing is actively regulated during cortical interneuron development

The extent to which AS is differentially regulated in cIN subtypes has not been explored. To address this question, we examined the amount of differentially expressed exons in cINs across developmental time-points ranging from embryogenesis (Embryonic day 18.5, E18.5) to juvenile ages (Postnatal day 22, P22). We took advantage of the Tg-*Lhx6e::GFP* line, in which green fluorescent protein (GFP) is constitutively expressed in medial ganglionic eminence (MGE)-derived cINs (PV+ and SST+ INs) at all ages (Figure 1A-B). Using fluorescence-activated cell sorting (FACS), we isolated GFP+ MGE-cINs at E18.5, P4, P8, P12 and P22. Sorted cINs were used to prepare cDNA libraries that were subsequently sequenced in order to investigate changes in the prevalence of alternatively spliced exons (spliced exon: SE, mutually exclusive exons: MEX, retained intron: RI, alternative 5’ splice site: A5SS, alternative 3’ splice site: A3SS; Figure 1C), (RNA-seq metadata in Supplementary Table 1). We next examined the pool of genes for which differentially expressed exons or isoforms were observed. Using rMATS (Shen et al., 2014), we found that the number of differentially expressed exons in the *Lhx6+* population greatly varies throughout development (E18.5 vs P4: 1109 exons; P4 vs P8: 915 exons; P8 vs P12: 651 exons; P12 vs P22: 772 exons) (Figure 1D). Notably, while some of the genes undergoing AS were observed to be spliced at all developmental time-points, others show specific developmental profiles (E18.5 vs P4: 49% (414/846 genes); P4 vs P8: 39% (273/705 genes); P8 vs P12: 31% (163/530 genes); P12 vs P22: 37% (222/602 genes). Interestingly, despite changes in gene expression, there were no notable differences in the proportion of the types of splicing events at different developmental time-points (i.e. spliced exon: SE, mutually exclusive exons: MEX, retained intron: RI, alternative 5’ splice site: A5SS, alternative 3’ splice site: A3SS) (Figure 1E-F; Supplementary Figure 1A-D).

**Figure 1:**
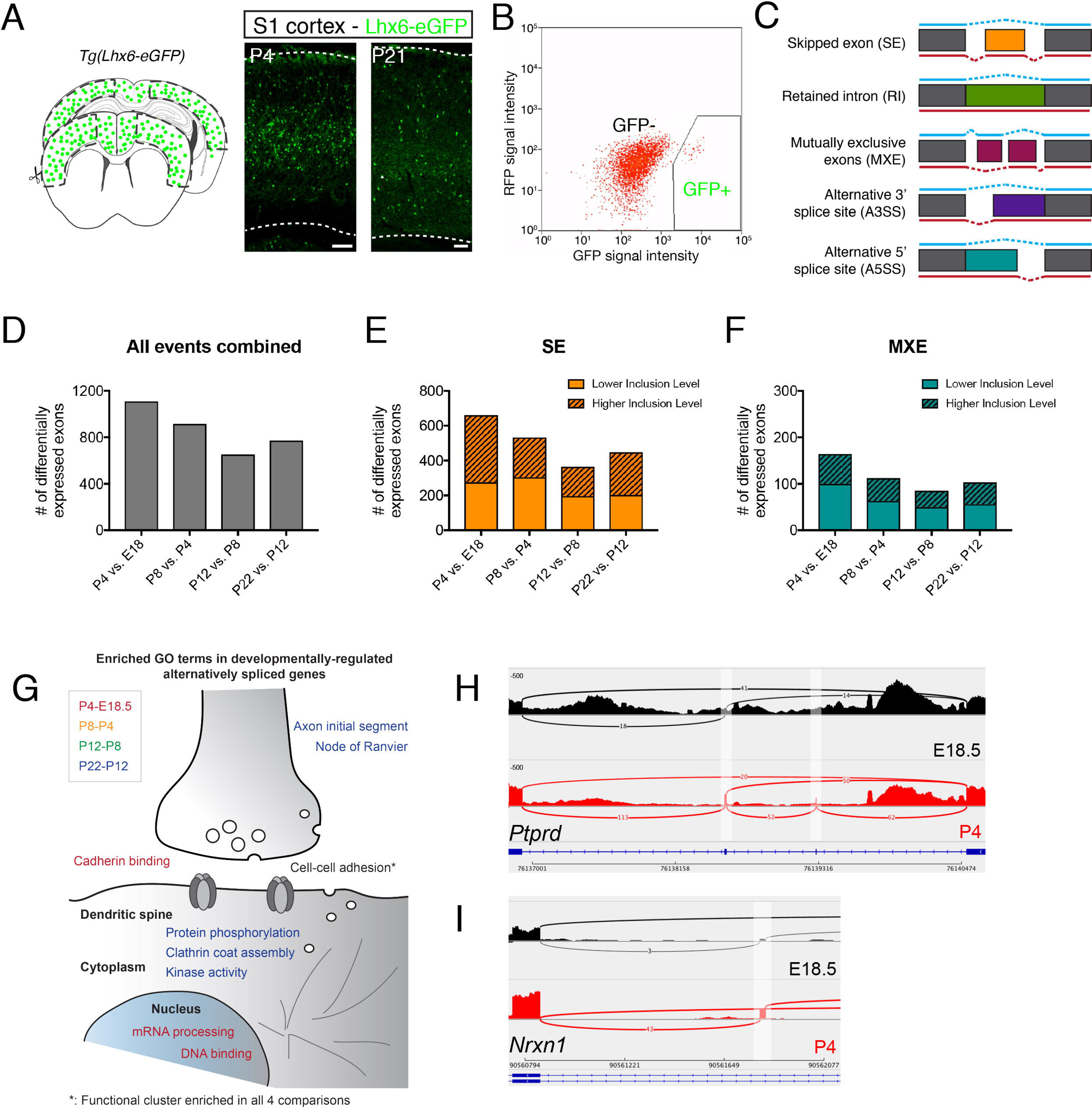
Alternative splicing is dynamically regulated during MGE cIN development. (A) Schematic depicting the cortical region isolated from P21 transgenic mouse brains. Corresponding regions were dissected from E18.5, P4, P8 and P12 brains. Examples of the MGE cINs revealed by immunostaining from P4 and P21 *Lhx6::eGFP* primary somatosensory cortices (S1) with anti-GFP (green) antibody. (B) Fluorescence-activated cell sorting of GFP+ interneurons from *Lhx6::eGFP* mouse cerebral cortex. (C) Schematic representing the five alternative splicing events detected by rMATS. (D) Left: Histogram depicting the number of differentially-expressed exons in the following comparisons: P4 vs. E18.5, P8 vs. P4, P12 vs. P8 and P22 vs P12 (rMATS analysis, *p*-val<0.05 and |∆ψ|≥0.1). (E-F) Histograms depicting the number of SE: spliced exons (E) and MXE: mutually-exclusive exons (F) that are either more excluded (skipped) (plain color) or included (spliced) (patterned-color). (G) Schematic depicting the biological functions of the enriched GO terms in the 4 following comparisons: P4 vs. E18.5 (dark red) and P22 vs P12 (blue). (H,I) Sashimi plots of representative examples of the differentially spliced transcripts *Ptprd* (H) and *Nrxn1* (I) from the P4 (red) vs. E18.5 (black) comparisons. Scale bars: (A) 100µm.

We next examined whether certain gene categories are more likely to be alternatively spliced at specific time-points by looking for enrichment of gene ontology (GO) terms across different developmental windows. Analysis between E18.5 and P4 revealed an enrichment of GO terms related to cell-cell adhesion, cadherin binding, mRNA processing and DNA binding (Figure 1G and Supplementary Table 2). At later developmental time-points (P12 vs P22), the GO enrichment analysis revealed active splicing of genes associated with the following categories: cell-cell adhesion, clathrin coat assembly, protein phosphorylation, kinase activity and node of Ranvier / Axon initial segment (Figure 1F, Supplementary Figure 1 and Supplementary Table 2). A commonality across all ages is the enrichment for AS genes related to synapse formation and cell-cell adhesion. AS events between E18.5 to P4 appear to be related to initial synaptic establishment and postsynaptic specialization (ex. *Ptprz1*, *Clasp1*, *Nrnx1*, *Ptprd*) (Figure 1H,I), whereas those enriched between P4 and P8 are involved in axon formation and presynaptic function (ex. *Ntng1*, *Ctnnd1*, *Ank2, Cacna1a, Erc2*). Finally, AS events between P12 and P22 are related to protein phosphorylation (ex. *Cask*, *Limk2*, *Map4k4*) and endocytosis (ex. *Picam*, *Sh3kbp1*), (Figure 1G and Supplementary Table 2).

### Rbfox1 localization in both PV+ and SST+ cortical interneurons varies during development

To identify specific factors mediating these AS events, we next surveyed known splice regulators for their expression in cINs. We identified that the RNA-binding protein, fox-1 homolog 1 (Rbfox1) is expressed in post-mitotic precursor cells as early as E14.5 (Figure 2A), a time at which cINs are migrating towards the cortex. Using immunofluorescence and genetic fate-mapping, we examined the expression of Rbfox1 within specific cIN subtypes. We observed that the number of Rbfox1-expressing cINs steadily increases during embryonic development up until juvenile stages. By P22 it is expressed in 68.6% (±9.5%) of *Dlx6a^cre^* fate-mapped cINs within the somatosensory cortex (Figure 2B,C). Notably, its expression is enriched in MGE-derived cINs (PV+ cINs: 100%; SST+ cINs: 73.7±9.5%; Figure 2D,E), with relatively fewer caudal ganglion eminence (CGE)-derived cINs expressing Rbfox1 (*5HT3aR::eGFP*: 27.5%±4.3%; Figure 2F).

**Figure 2:**
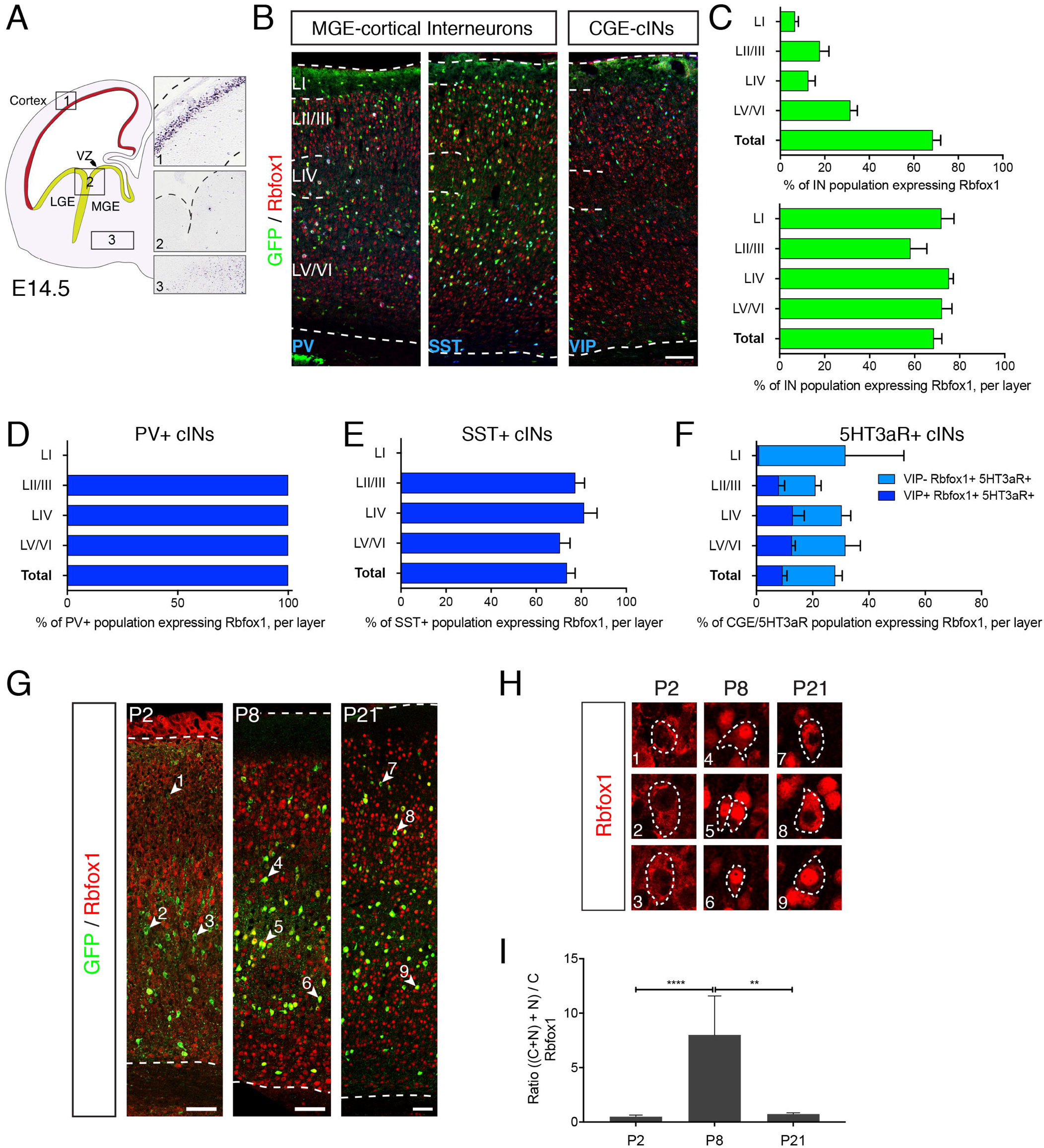
Within cortical interneurons, the splicing regulator Rbfox1 is preferentially expressed in the PV and SST expressing subtypes. (A) *In situ* hybridization on a E14.5 mouse embryo (GenePaint.org) revealing that the expression of *Rbfox1* is restricted to postmitotic neurons within the cortical plate and is excluded from the ventricular zones of the dorsal cortex (1) and ganglionic eminences (2). In addition, sparse Rbfox1-expressing neurons can be found in the mantle of the ganglionic eminences (3). (B) Immunostaining of P21 *Dlx6a^Cre^;RCE^eGFP/+^* (cortical interneurons) and *5HT3aR::eGFP* (CGE cINs) S1 cortices using anti-GFP (green), anti-Rbfox1 (red) and anti-PV (blue, left), anti-SST (blue, middle) and anti-VIP (blue, right). (C) Top: Histogram depicting the percentage of the total cIN population (fate-mapped using *Dlx6a^Cre^;RCE^eGFP/+^*) that express Rbfox1 within cortical layers (Layer I:LI, Layers II/III: LII/III, Layer IV: LIV and Layers V/VI: LV/VI) and in the whole cortex (Total). Bottom: Histogram depicting the percentage of the cIN population (fate-mapped using *Dlx6a^Cre^;RCE^eGFP/+^*) per layer that express Rbfox1 and in the whole cortex (Total). (D) Histogram depicting the percentage of the PV+ expressing cIN population (fate-mapped using *Dlx6a^Cre^;RCE^eGFP/+^* and stained with anti-PV) that express Rbfox1 in various layers and in the whole cortex (Total). (E) Histogram depicting the percentage of the SST+ expressing cIN population (fate-mapped using *Dlx6a^Cre^;RCE^eGFP/+^* and stained with anti-SST) that express Rbfox1 in various layers and in the whole cortex (Total). (F) Histogram depicting the percentage of the *5HT3aR::eGFP* cIN population (stained with anti-VIP) divided between eGFP/VIP+ (yellow) and eGFP/VIP- (brown) that express Rbfox1 in various layers and in the whole cortex (Total). (G) Immunostaining of P2, P8 and P21 S1 cortex from *Lhx6::eGFP* transgenic mice, where MGE-derived cINs are labelled with GFP and stained with anti-Rbfox1 (red). Nine representative GFP+ cINs are labelled with white arrowheads (1-9) and the corresponding higher magnifications pictures are shown in B. (H) Higher magnifications illustrate a shift in Rbfox1 localization between P2-P8 and P8-P21 (anti-Rbfox1, red). (I) The ratios of nuclear Rbfox1 (Rbfox1_N & Rbfox1_C+N) over cytoplasmic Rbfox1 (Rbfox1_C) at P2, P8 and P21 in eGFP+ interneurons were determined based on the quantifications shown in Suppl. Figure 2E (N_P2_=16; N_P8_=6; N_P21_=6). Scale bars: (B) 100µm; (G) 60µm; (H) 15µm; insets: 20µm.

The *Rbfox1* gene contains several promoters and exons that can result in the generation of multiple isoforms. The nuclear versus cytoplasmic localization of Rbfox1 is controlled by the skipping or inclusion of Rbfox1 mRNA’s exon 19. The skipping of exon 19 unveils a cryptic nuclear localization signal (NLS) in downstream exon 20 (Lee et al., 2009). It is this isoform that functions as a splicing regulator (Jin et al., 2003; Nakahata and Kawamoto, 2005; Underwood et al., 2005). Conversely, the cytoplasmic isoform of Rbfox1 (exon 19+) has recently been shown to regulate the stability and enhance the expression level of its target mRNAs (Lee et al., 2016). Given our observation that the number of differentially expressed exons varies during MGE-cIN development, we examined whether Rbfox1 localization also changes. We quantified the relative proportion of *Lhx6+* MGE-cINs in which Rbfox1 was enriched within the cytoplasm (Rbfox1_C), the nucleus (Rbfox1_N), or in both the nucleus and cytoplasm (Rbfox1_C+N) (Supplementary Figure 1H). We found that at P2, 67.3% (±6.5) of *Lhx6::eGFP+* cINs express Rbfox1_C, 27.3%±5.3 express Rbfox1_C+N and only 5.5% (±3.9) express Rbfox1_N (Figure 2G,H, Supplementary Figure 1I). Interestingly at P8, we observed a significant increase of the proportion of Rbfox1_C+N (86.8%±7.8) at the expense of Rbfox1_C, whose proportion decreases to 6.50% (±7.5) (Supplementary Figure 1I). As a consequence of this change in distribution, the ratio of neurons with nuclear Rbfox1 (including Rbfox1_C+N) to neurons expressing Rbfox1_C becomes significantly higher (8.00 vs. 0.50) (Figure 2I). Interestingly, the reduction in Rbfox1_C is transient, as indicated by the enrichment of Rbfox1_C (58.2%±4.0) at P21, which lead to a significant reduction of the N/C ratio (0.73 vs. 8.00) (Figure 2I, Supplementary Figure 1I). Altogether, our observations indicate that at P8 Rbfox1 may be primarily involved in regulating splicing. By contrast, the later enrichment of Rbfox1_C suggests that in more mature cells (P21) Rbfox1 may additionally regulate mRNA stability (Vuong, 2018). Notably, this trend is similar within both SST+ (*Lhx6::eGFP+/SST+)* and PV+ (*Lhx6::eGFP+*/SST) cINs over this time period (not shown).

### Interneuron-specific loss of Rbfox1 impairs cortical inhibition

We next conditionally inactivated *Rbfox1* by crossing the *Rbfox1^F/F^* allele (Gehman et al., 2011) with a *Dlx6a^Cre^* driver line (Yu et al., 2011). This results in the deletion of *Rbfox1* in all cINs during embryogenesis, shortly after they become postmitotic. We took advantage of two different reporter lines, the *RCE^LoxP^* (referred to as *RCE^eGFP^*) and *Ai9^LoxP^* (referred to as *Ai9^F/F^*), which upon Cre-mediated recombination conditionally express GFP or TdTomato, respectively. We confirmed by immunofluorescence that Rbfox1 expression is abolished in cINs in the *Dlx6a^cre^* conditional *knock-out* (*Dlx6a^Cre^;Rbfox1^F/F^;RCE^eGFP^*: henceforth referred to as *Dlx-cKO*, Supplementary Figure 2A). We observed that *wild-type (wt)*, *Dlx-het* and *Dlx-cKO* are born at the expected mendelian ratios (1:2:1; not shown). However, subsequently we observed an increased lethality of *Dlx-cKO* mice beginning at P18, while both *wt* and *Dlx-het* show normal survival (Supplementary Figure 2B). All *Dlx-cKO* mice were deceased by P45. In several instances, we observed *Dlx-cKOs* spontaneously experiencing seizures in their home cage, suggesting that the shortened lifespan observed was caused by lethal seizures. To determine whether *Dlx-cKOs* exhibited increased seizure susceptibility, we monitored the behavior of P12-P14 *wt* and *Dlx-cKO* mice following intraperitoneal injections of kainic acid (KA, 12mg/kg) and scored their behavior over a 70 min period according to a modified Racine Scale (Gehman et al., 2011; Racine, 1972). We found that KA elicited *status epilepticus* and death in *Dlx-cKO* mice within 30 min, while *wt* mice experienced mild seizures (2/9 mice displayed forelimb clonus) and eventually recovered (Supplementary Figure 2C). Notably, heightened seizure susceptibility similar to what we observed in *Dlx-cKO* mice has been reported in adult mice following the pan-neuronal removal of *Rbfox1* using the *Nestin^Cre^* (Gehman et al., 2011). This suggests that both the pan-neuronal and the cIN-specific loss of *Rbfox1* heightens seizure susceptibility and potentially impairs inhibition. However, the interneuron-specific model is more severely affected as suggested by the early lethality observed in *Dlx-cKO* mice.

The literature abounds with examples where reductions in cIN numbers result in seizures (Lodato et al., 2011; Marín, 2012). However, we did not find a reduction in cINs. To the contrary, we found a slight increase in PV+ and SST+ cIN density suggesting a decrease in cell death (Priya et al., 2018) (Supplementary Figure 2D-G). The seizure susceptibility in *Dlx-cKO* mice in the absence of a loss of interneurons suggested that inhibitory function was compromised in these animals. To examine this issue in greater detail, we assessed inhibitory currents in the somatosensory cortex using whole-cell patch-clamp recording in acute slices of both juvenile *Rbfox1 Dlx-cKO* and *Dlx-Control* littermates (aged P15-P18). We recorded and analyzed the level of inhibition received by pyramidal neurons in LII/III. We observed a significant reduction in both the frequency (3.12±0.53 Hz in *Dlx-cKO* vs 4.86±0.42 Hz in *Dlx-Control*) and amplitude (median amplitude: 13.10 pA in *Dlx-cKO* vs 14.72 pA in *Dlx-Control*) of miniature inhibitory post-synaptic currents (mIPSCs) in the mutant brains (Figure 3A,B). Consistent with a loss of inhibition and seizure activity, we detected a substantial increase in the expression of activity-dependent IEG cFos (Figure 3C,D) and the neuropeptide NPY (not shown) in all cells. In sum, *Dlx-*cKO mice display impairments in inhibitory function in the cortex that likely explain the increased seizure susceptibility.

**Figure 3:**
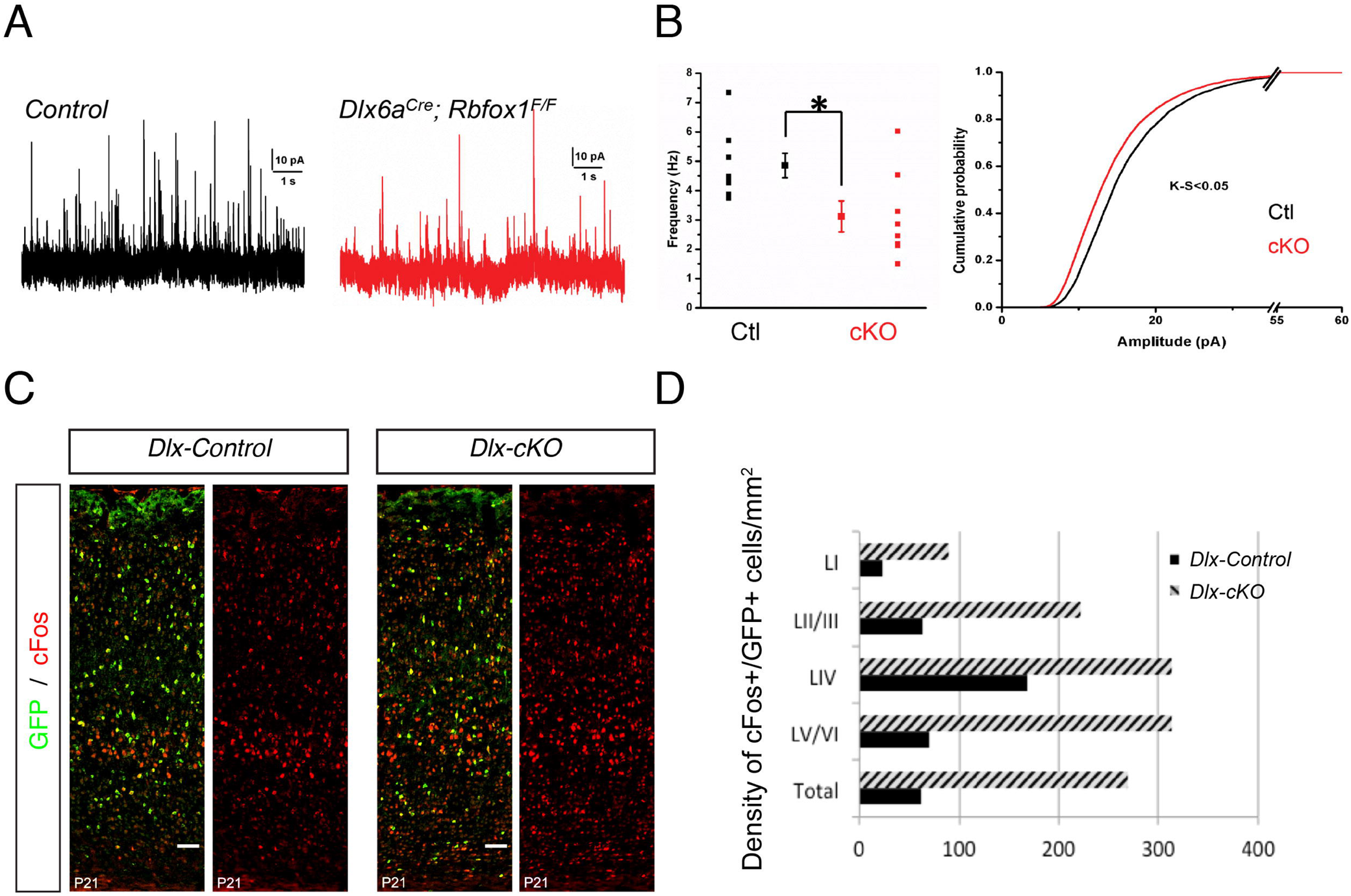
The interneuron-specific inactivation of *Rbfox1* impairs inhibitory function within the cortex. (A) Representative mIPSCs recorded from LII/III pyramidal neurons in control (*Dlx-Control*; black) and *Dlx-cKO* (red) cortex at P17-P18. (B) Left: Averaged mIPSC frequency between *Dlx-Control* and *Dlx-cKO* is reduced (N*_Dlx-cKO_*=8 and N*_Control_*=8; *p-*val=0.02). Right: Cumulative probability of mIPSC amplitudes from the recorded pyramidal excitatory neurons is also reduced (Kolmogorov-Smirnov test, *p*-val<0.05). (C) Immunostaining of S1 cortex at P21 using anti-GFP (green) and anti-cFos (red) antibodies in the *Dlx-control* and *Dlx-cKO* backgrounds. (D) Associated quantification of cFos+/GFP+ interneurons across cortical layers. Scale bars: (C) 100µm.

### *Rbfox1* inactivation in SST+ and PV+ cortical interneurons impairs their efferent connectivity

Epileptic phenotypes have previously been reported to be due to altered maturation of the intrinsic properties of MGE-derived cINs (Batista-Brito et al., 2009; Close et al., 2012). In order to test whether the physiological membrane properties of MGE cINs are affected by the cell autonomous removal of *Rbfox1*, we generated SST+ and PV+ interneuron specific *Rbfox1* conditional mutants using *SST^Cre^* (*SST-cKOs*) and *Tac1^Cre^* (*Tac1-cKOs*) drivers, respectively (Harris et al., 2014; Taniguchi et al., 2011). The *Tac1^Cre^* driver allows the targeting of a subset of LII/III (46.5±11.1%) and LV/VI (52.3±10.3 %) PV+ cINs earlier than the standard *PV^Cre^* drivers (Supplementary Figure 3A,B). As 24% of *Tac1^Cre^-*targeted cells are not cINs (likely pyramidal neurons, not shown), we used an intersectional genetic strategy to confine our analysis to PV+ cINs. We employed the *Dlx5/6^FlpE^* allele, which drives the expression of the FlpE recombinase in all cINs, and the *Tac1^Cre^* driver allele in combination with the intersectional TdTomato-reporter Ai65. Using this approach, we first examined their firing (first spike characteristics and maximum firing frequency, rheobase) and passive membrane properties (input resistance) using targeted whole-cell patch-clamp recordings in acute slices of juvenile (aged P15-P18) *SST- or Tac1-cKO* versus *control* littermates (*Tac1-cKO*: 8 HET and 13 MUT; *SST-cKO*: 13 HET and 15 MUT). Most parameters were unaffected in SST-conditional *Rbfox1* mutants, with the exception of an increased maximum firing frequency in *SST-cKO*s as compared to *SST-Controls* (107±7 Hz vs 85±7 Hz in *SST-cKO* vs *SST-control*, Supplementary Table 3). The examination of *Tac1-cKO* cINs did not reveal any significant alteration of their active or passive electrophysiological properties (Supplementary Table 3).

As we did not identify major defects within the intrinsic properties of PV+ and SST+ cINs, we explored whether the loss of inhibitory drive in *Dlx-cKO* mutants reflects defects in the synaptic connectivity of these populations (Figure 4A). We used conditional genetic reporters and cell-type specific synaptic markers to determine whether efferent synapses from PV+, SST+ cINs or both were affected. To examine this question within PV+ cINs, we took advantage of the PV+ IN-specific expression of the presynaptic protein Synaptotagmin-2 (Syt2) (Sommeijer and Levelt, 2012) for analyzing PV+ axon terminals in *Dlx-cKO* at weaning age (P19-P23). Using immunohistochemistry, confocal imaging and puncta analysis (Ippolito and Eroglu, 2010), we quantified the number of Syt2+ puncta that co-localized with the postsynaptic marker Gephyrin within perisomatic regions of LII/III and LV/VI pyramidal neurons. We observed that the removal of *Rbfox1* caused a reduction in the number of PV-specific synapses comparing controls to mutants (0.343±0.012 A.U. and 0.216±0.008 A.U., respectively) (Figure 4B-C; Supplementary Figure 3C). Hence, the reduction of PV-specific inhibitory synapses likely accounts at least in part for the reduced mIPSC frequency.

**Figure 4:**
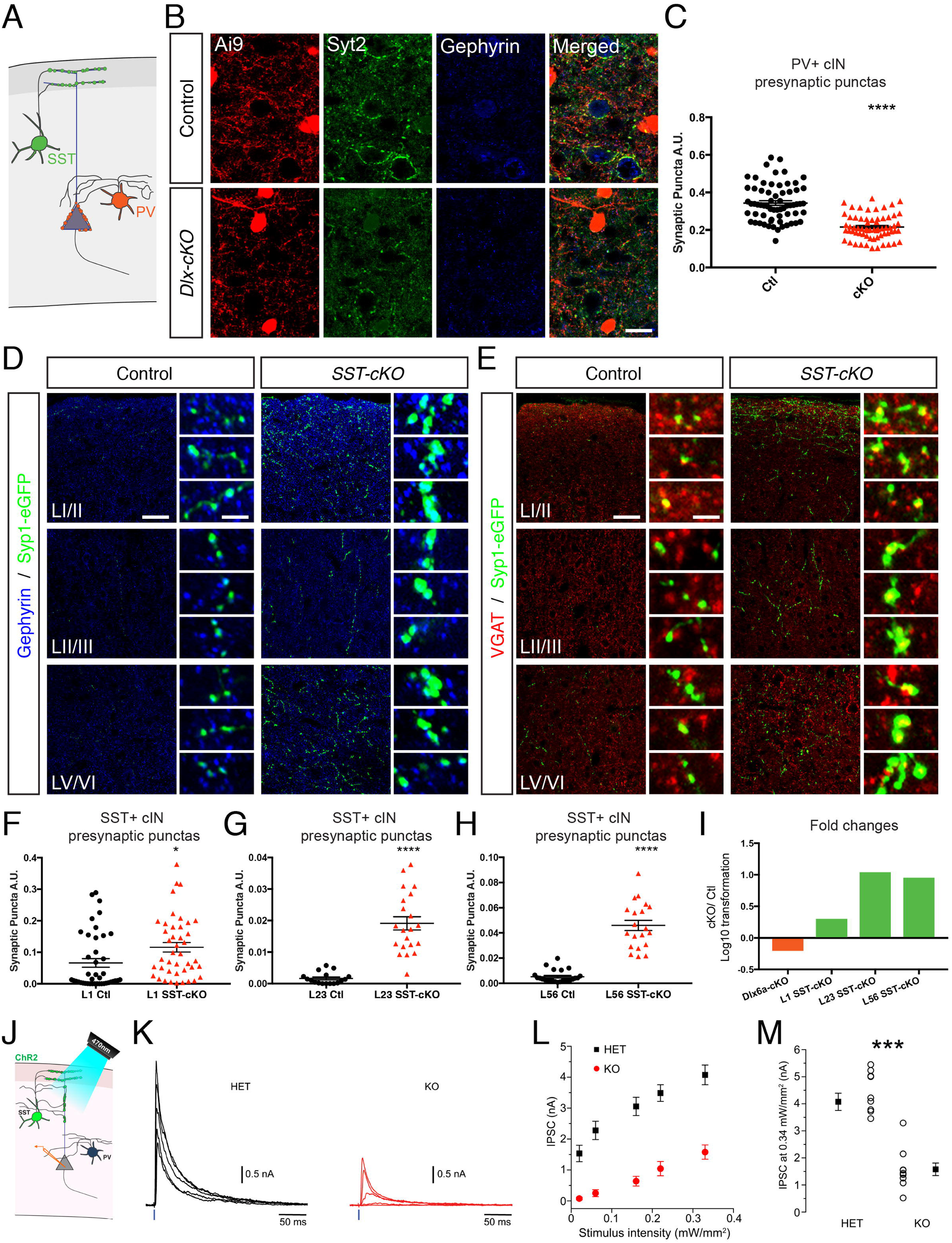
Inactivation of Rbfox1 has opposing effects on somatostatin+ and parvalbumin+ cIN efferent connectivity. (A) Schematic illustrating the subcellular targeting of SST+ cIN efferent synapses on apical dendrites of pyramidal neurons (green) and of PV+ cIN efferent perisomatic synapses onto pyramidal neurons (orange). (B) Immunostaining of *Dlx-Control* and *Dlx-cKO* cortex with anti-dsRed (red), anti-Syt2 (green) and anti-Gephyrin (blue). (C) Quantification of Ai9/Syt2/Gephyrin puncta in both control and Dlx-cKO at P17-P18 reveals a reduction in PV+ cIN efferents (N*_Dlx-cKO_*=65, N*_Dlx-Control_*=65). (D) Immunostaining of *SST-Control* and *SST-cKO* cortex using anti-GFP (green) and anti-Gephyrin (blue) in Layer I/II (LI/II), Layer II/III (LII/III) and Layer V/VI (LV/V). This staining reveals the presence of Syp-eGFP synaptic reporter fusion protein colocalized or in close apposition with the postsynaptic reporter Gephyrin. (E) Immunostaining of *SST-Control* and *SST-cKO* cortex with anti-GFP and anti-VGAT using anti-GFP (green), anti-VGAT (red) in LI/II, LII/III and LV/V. This staining reveals the presence of Syp-eGFP synaptic reporter fusion protein colocalized or in close apposition with the presynaptic reporter VGAT. (F) Quantification of GFP/Gephyrin puncta in both *Controls* and *SST-cKOs* at P21 in Layer I/II (N_Control_=N_SSTcKO_=42), (G) in Layer II/III (N_Control_=N_SSTcKO_=21) and (H) in Layer V/VI (N_Control_=N_SSTcKO_=24). (I) Log10 transformations of the synaptic puncta fold-changes observed between cKO and controls and shown in (C) and (F-H). (J) Illustration of the SST+ cIN targeted optogenetics experiment: ChR2 is expressed in SST+ cINs using the Ai32 reporter line. SST+ cIN neurotransmission was activated with 470nm light stimulation and the inhibitory response was recorded by whole-cell patch clamp recordings from layer II-III pyramidal neurons within the somatosensory cortex. (K) Representative evoked currents from a control (black trace) and a cKO (red trace) mice. IPSCs were evoked by a series of 1 ms steps of increasing light intensity. (L) Input-output curve representing the amplitude of the IPSC (nA) as a function of the light stimulus intensity (mW/mm^2^). (M) Summary plot of the response to the maximal light intensity. Scale bars: (B) 25µm, (D,E) 60µm, insets 5µm.

We next tested whether a later removal of *Rbfox1* results in a similar phenotype using *PV^cre^* driver mice. Expression of the *Pvalb* gene begins around P14 in the somatosensory cortex, resulting in *Rbfox1* inactivation during the third postnatal week, a period following peak synaptogenesis. Notably, we found that the late removal of *Rbfox1* in PV+ cINs did not result in a decrease in the density of Syt2+/Gephyrin+ co-localized puncta (Supplementary Figure 4A) or PV+ cINs (Supplementary Figure 5B). These results support our hypothesis that Rbfox1 is critically required during the first two postnatal weeks, coincident with the period of cortical synaptogenesis.

To investigate the consequences of *Rbfox1* genetic inactivation on SST+ cINs efferent synapses, we crossed compound *SST^cre^;Rbfox1^F/F^* mice onto a background carrying the conditional presynaptic reporter *Syp-eGFP* (Li et al., 2010). In combination with anti-Gephyrin immunostaining, we were able to assess SST+ cortical interneurons’ efferent inhibitory synapses. We quantified the number of GFP+/Gephyrin+ puncta using confocal imaging in P20-P25 S1 mouse cortex. To our surprise, we observed a significant increase in the amount of SST-cIN synaptic puncta across all layers of the cortex in *SST-cKOs* as compared to control littermates (Figure 4D,F-H) (L1: 0.066±0.014 vs. 0.116±0.015 and L2/3: 0.002±0.001 vs. 0.019±0.002, in *SST-Control* vs *SST-cKO*, respectively). In particular, we found many more inhibitory synapses in the deeper cortical layers (LV/VI) of *SST-cKO* than in control animals (L5/6: 0.005±0.001 vs. 0.046±0.004, in *SST-Control* vs *SST-cKO*). To further ascertain that we are accurately assessing the increase in synapse number *in SST-cKO* animals, we confirmed the juxtaposition of Syp-GFP to both VGAT (vesicular GABA transporter) (Figure 4E; Supplementary Figure 3D-F) or Synaptophysin-1 protein using immunocytochemistry (not shown). Altogether, our investigations uncovered distinct but opposing synaptic alterations in PV- versus *SST-cKO* mutants (Figure 4I). Upon the conditional removal of *Rbfox1*, PV+ cINs showed a reduced number of efferent synapses whereas SST+ cINs form supernumerary inhibitory efferent synapses.

### Conditional loss of *Rbfox1* reduces efferent synaptic transmission from SST+ cortical interneurons

To reconcile the increased density of efferent synapses in *SST-cKO* mutants with the observed reduction in inhibitory drive, we examined whether the extra synapses we observed were functional. In order to assess the synaptic output from SST-cINs onto pyramidal neurons, we employed an optogenetic-based approach. Channelrhodopsin2 (ChR2) was expressed in SST+ interneurons using the Ai32 conditional ChR2-expressing reporter (*Ai32^F/F^*). We applied single 1ms blue-light stimulations (470nm) of varying intensities onto layer 2/3 of somatosensory cortical slices and analyzed averaged resultant IPSCs within pyramidal neurons. We observed a dramatic reduction of inhibitory neurotransmission from SST+ cINs in *SST*-*cKO* adult slices (Figure 4J-M) (4.1±0.3 nA vs. 1.6±0.2 nA, *p*-val=5.35E-6). To determine whether the increased number of efferent SST+ cIN synapses was developmental or compensatory in nature, we analyzed the quantity of synaptic puncta at an earlier age (P8) and found that the number of inhibitory puncta was unchanged (Supplementary Figure 4C,D). This suggested that the supernumerary SST+ synapses represent a homeostatic compensation resulting from the lack of functional SST cIN axon terminals.

### Rbfox1 mediates alternative splicing of transcripts encoding presynaptic proteins

To investigate alterations in gene expression and AS among transcripts within PV and SST cIN subtypes, we selectively sorted these populations from *Rbfox1* conditional mutant and control animals at P8, a developmental stage that corresponds to the peak of synaptogenesis. This also represents a time-point when Rbfox1 is predominantly localized within the nucleus of these cINs. We first examined whether loss of *Rbfox1* affected gene expression. In SST-cKOs we identified the up-regulation and the down-regulation of only 2 and 4 genes, respectively (Supplementary Table 4; |Log2FC|>1; p-value<0.01). By contrast, the *Tac1*-inactivation of *Rbfox1* caused substantial changes in gene expression. We observed 11 up- and 192 down-regulated genes, respectively (Supplementary Table 4; |Log2FC|>1; *p*-val<0.01).

We next examined how the selective loss of *Rbfox1* in *SST-cKO* and *Tac1-cKO* affected AS. Using rMATS analysis, we identified 305 exons within SST+ cINs that where alternatively spliced due to the inactivation of *Rbfox1* (Figure 5A,B; Supplementary Table 6). GO analysis carried out on the 278 genes displaying one or more Rbfox1-dependent splicing changes revealed a significant enrichment of genes with synaptic function (Figure 5C,E, Supplementary Table 5). Similar analysis of the *Tac1*-specific inactivation of *Rbfox1* affected the splicing of 742 alternatively spliced transcripts (Figure 5A,B, Supplementary Table 6). In this case, GO term enrichment analysis revealed 580 differentially spliced genes that were preferentially associated with cell-cell adhesion (Figure 5D, E, Supplementary Table 5).

**Figure 5:**
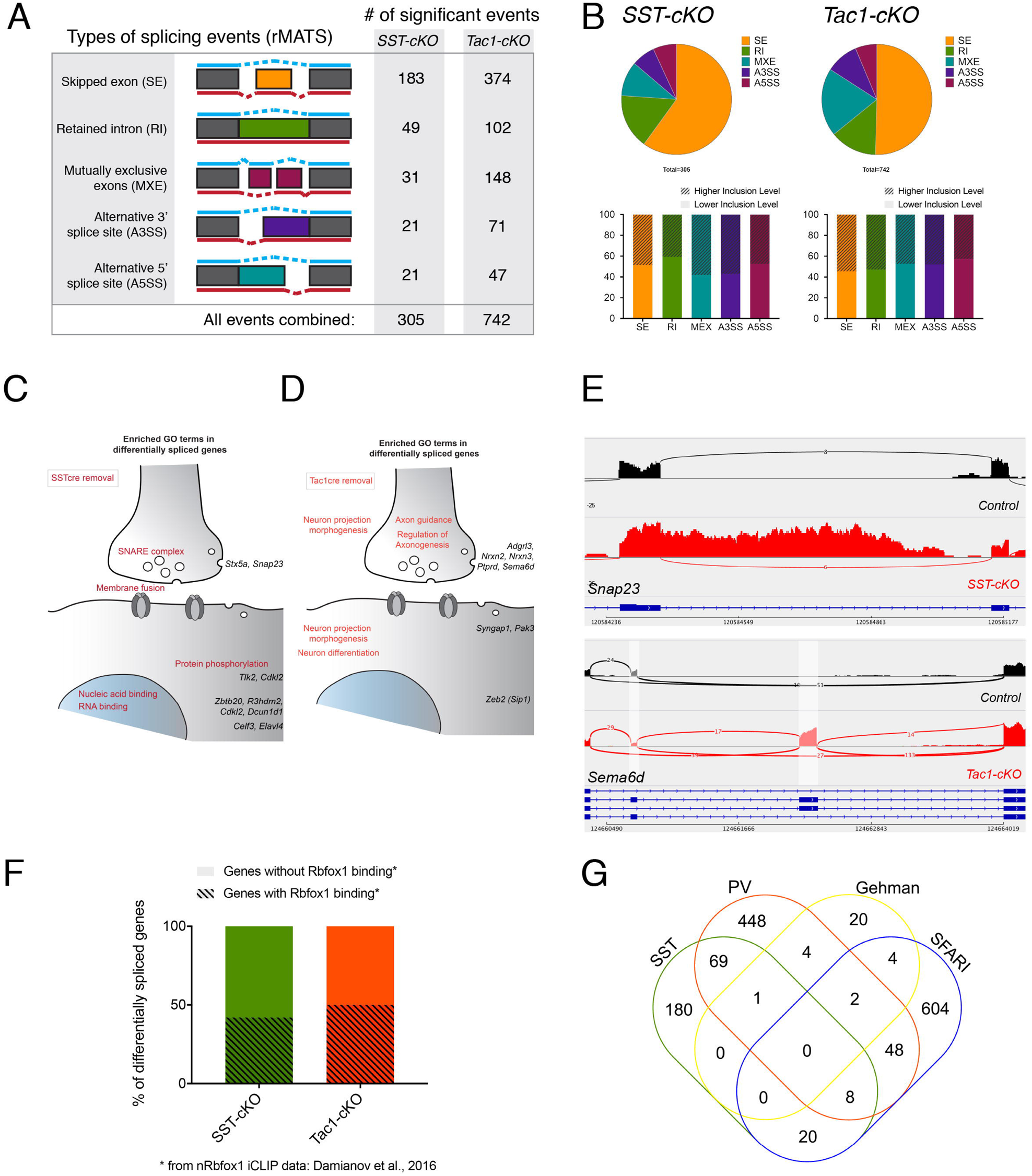
Alternative splicing programs in developing SST+ and PV+ cortical interneurons are altered upon loss of *Rbfox1*. (A) Schematic representation of the five types of alternative spliced exon events detected by rMATS and the number of statistically significant events (*p-val* <0.05, |∆ψ| >0.1) detected within the *SST-cKO* and the *Tac1-cKO* as compared to their respective controls at P8. (B) Top: pie charts showing the break-down of AS events in the *SST-cKO* (left) and *Tac1-cKO* (right) relative to their respective controls at P8 (*p-val* <0.05, |∆ψ| >0.1). Bottom: Proportion of alternative splicing events of excluded (plain color) or included target exons (patterned-color) in either comparisons. (C,D) Schematics depicting the different biological functions of the enriched GO terms from the set of differentially spliced genes in P8 *SST-cKO* (C) and P8 *Tac1-cKO* cINs (D). (E) Sashimi plots of representative examples of differentially spliced transcripts in P8 *SST-cKO* (*Snap23*) and P8 *Tac1-cKO* (*Sema6d*). (F) Relative fractions of the differentially spliced genes found in either cell type from *SST-cKOs* or *Tac1-cKOs* with or without evidence of nuclear Rbfox1 direct binding as identified by iCLIP experiment from a nuclear fraction of forebrain neurons (Damianov et al., 2016). (G) Venn diagram showing the intersection between the sets of differentially spliced transcripts identified in either cell types from *SST*- and *Tac1*-*cKOs* and the data set of SFARI ASD-candidate genes and the previously identified set of 30 genes differentially spliced in the adult brain of *Nestin-Cre Rbfox1* conditional KOs (Gehman et al., 2011).

We observed that a large proportion of alternatively spliced transcripts within SST+ and PV+ cINS – 58% (162/278) and 50% (288/580), respectively – had not been previously identified as direct nuclear Rbfox1 targets in the adult forebrain (Figure 5F, Supplementary Figure 6) (Damianov et al., 2016). It is possible that some of the identified transcripts without reported Rbfox1 direct binding might be differentially spliced only as an indirect consequence of *Rbfox1* inactivation. Notably, we found that transcripts coding for other *trans*-acting RNA splicing regulators, such as Elavl4 (Figure 5C, Supplementary Figure 5), were differentially spliced as a result of *Rbfox1* inactivation. Thus, it is likely that the loss of *Rbfox1* also indirectly alters AS patterns indirectly by impacting Elavl4 function. Of note, previous work examining the pan-neuronal (i.e. *Nestin^cre^*) removal of *Rbfox1* (Gehman et al., 2011) identified 31 transcripts whose AS was altered and of these only very few were identified within our analysis (1/31 in SST-cINs and 7/31 in PV-cINs) (Figure 5G). However, given the small proportion of cINs within the cortex, it remains a possibility that PV+ and SST+-specific *Rbfox1*-dependent events were simply undetected in previous investigations of whole cortical tissue.

In humans, *RBFOX1* is recognized to be associated with autism spectrum disorders (Martin et al., 2007; Sebat et al., 2007), epilepsy and intellectual disability (Bhalla et al., 2004). Given the enrichment of ASD-associated genes identified as RBFOX1/Rbfox1 targets (Voineagu et al., 2011; Weyn-Vanhentenryck et al., 2014), we investigated whether AS in PV+ and SST+ *cKO* mutants preferentially affected ASD candidates. We hence examined the level of overlap between our putative *Rbfox1* targets and the 686 murine orthologs of ASD candidate genes (SFARI gene 2.0) (Abrahams et al., 2013). We found that 10% (28/278, SST-cINs; and 58/580, PV-cINs) of the genes we identified in either cell type are known ASD candidate genes (Figure 5G). This level of enrichment is comparable to the enrichment previously reported using pan-neuronal genetic manipulations (Weyn-Vanhentenryck et al., 2014).

### Rbfox1 orchestrates cell-type specific splicing of exons in cortical interneurons

The conditional removal of *Rbfox1* within either SST+ or PV+ cINs resulted in a total of 1047 AS events (Figure 6A), 954 events of which were unique to one of the two cell types (Figure 6A). Among these, only a negligible number of events involved genes that were differentially expressed between SST+ and PV+ cINs (0/954 and 90/954 in SST+ and PV+ cINs, respectively). Importantly, this shows that the cell-specific changes in splicing observed upon *Rbfox1* loss of function are not merely a result of differences in gene expression. Of the 93 events that occurred within both SST+ or PV+ cINs, 36 represent exons that were similarly spliced (or skipped), while 57 reflected cell-type specific AS outcomes (Figure 6A).

**Figure 6:**
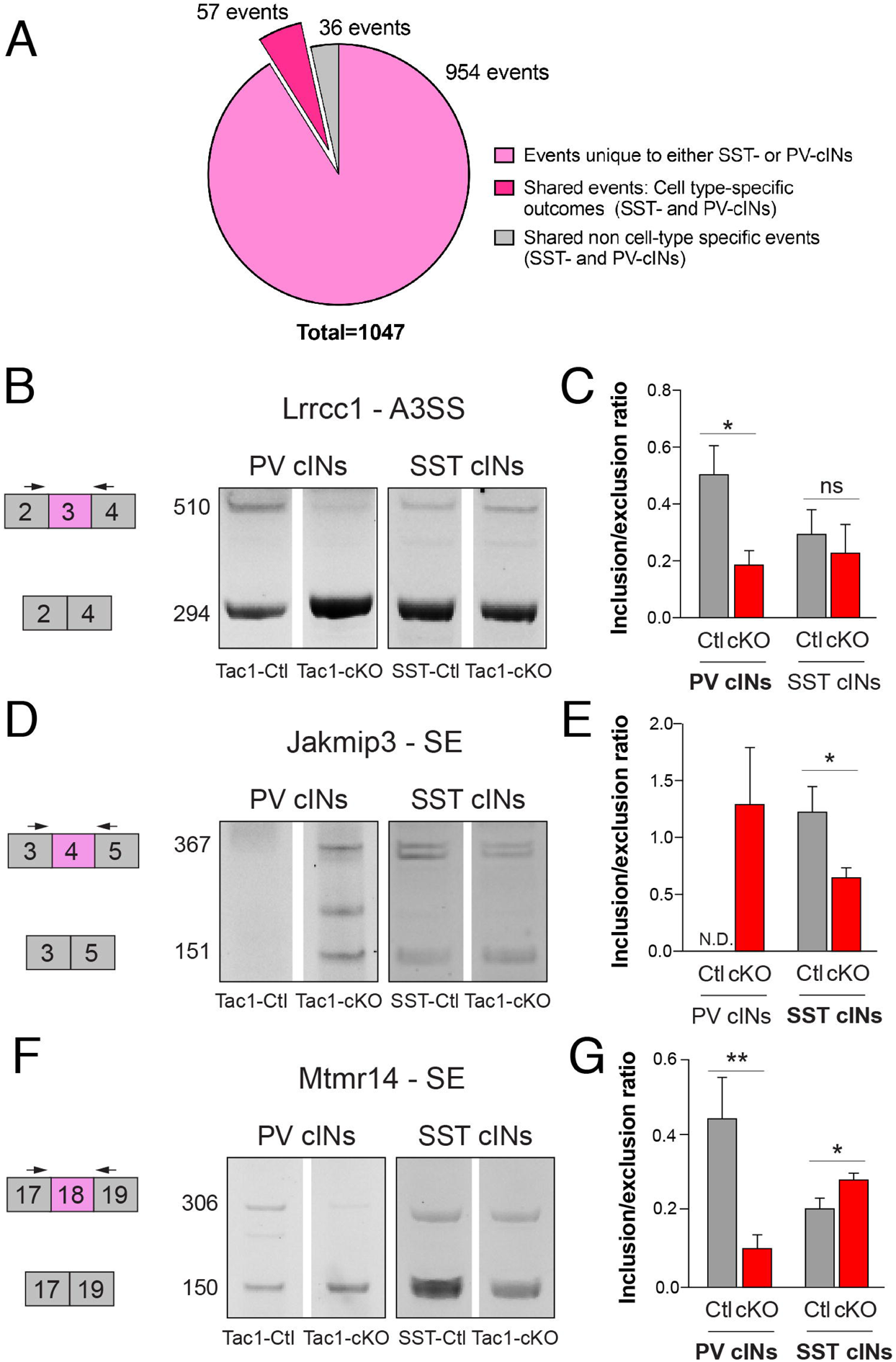
The splicing regulator Rbfox1 governs cell-type specific alternative programs in SST+ and PV+ cortical interneurons. (A) Pie chart showing the relative proportion of cell-type specific events (uniquely identified in either *SST*- or *Tac1-cKOs*, or in both *SST*- and *Tac1-cKOs* but with opposite outcomes) and non cell-type specific events. (B-G) Validation by fluorescent reverse transcription–polymerase chain reaction (RT-PCR) of the indicated Rbfox1 targets. Representative images (B, D, F) and quantification (C, E, G) showing the inclusion/exclusion ratio of Rbfox1-dependent exons in PV+ and SST+ controls versus *SST*- or *Tac1-*cKOs. N.D.: not detected. Data shown are averages ± SEM from ≥ 3 biological replicates.

We next sought to confirm that these AS events were altered in a cell-specific manner upon the conditional removal of *Rbfox1*. We therefore performed targeted validation experiments in sorted PV+ and SST+ cINs, using fluorescent RT-PCR amplification with primers flanking the alternatively spliced segments (Figure 6B-G and Figure S6E-H). The targets we validated included *Lrrcc1* (Leucine Rich Repeat And Coiled-Coil Centrosomal Protein 1), which encodes a docked synaptic vesicle component enriched at GABAergic synapses (Boyken et al., 2013). Although *Lrrcc1* is expressed in both PV+ and SST+ interneurons, we observed a lower inclusion of the spliced 3’ UTR exon in *Tac1-*but not *SST-cKO* mice (Figure 6B and 6C). Conversely, we found that the inclusion ratio of exon 4 in *Jakmip3* (Janus Kinase and Microtubule Interacting Protein 3) was reduced in *SST-* but not *Tac1-cKO* mice (Figure 6D and 6E). Perhaps most strikingly, the alternative splicing of exon 18 in *Mtmr14* (*Myotubularin Related Protein 14*) was differentially altered in the two cell types. Specifically, the absence of *Rbfox1* led to a decreased inclusion in *Tac1-cKO* mice but an increased inclusion in *SST-cKO* mice (Figure 6F and 6G). Altogether, these results confirmed that the changes in AS upon conditional loss of *Rbfox1* are markedly divergent between the two cell populations.

## Discussion

Our work highlights the surprising specificity of AS within particular cIN populations and during specific developmental events. In PV+ and SST+ cINs we observe striking changes in AS during the first three postnatal weeks and demonstrate that this process is both dynamic and exquisitely regulated. We discovered that this process is partially dependent upon Rbfox1, which has key cell-type specific roles during the maturation and integration of discrete cIN classes. Until recently AS had only been investigated within whole mouse brains or within heterogeneous cellular populations and investigated its role only at set stages (Dillman et al., 2013; Zhang et al., 2014). Recent work has however been able to implicate AS in neurogenesis and cell fate determination (Linares et al., 2015; Zhang et al., 2016), synaptic maintenance and plasticity (Iijima et al., 2011; Nguyen et al., 2016; and reviewed in Raj and Blencowe, 2015; Vuong et al., 2016). Our work expands upon these efforts by examining the cell type-specific consequences of AS during specific stages of cIN development. The robust changes in AS between E18.5 and P4 occur concomitantly with the settling of cINs within the cortical layers. This is a period during which they experience an increasingly dynamic range of neuronal activities and are robustly forming or losing synaptic contacts (Allene et al., 2008; Minlebaev et al., 2011; Yang et al., 2013; 2009). The concurrence of AS and activity during these stages of development coupled with the finding that activity alters Rbfox1 localization (Lee et al., 2009) warrants further investigation.

Our work has also clarified the requirement for *Rbfox1* for brain development. While the whole brain inactivation of *Rbfox1* causes seizures and results in lethality (Gehman et al., 2011), the neural and molecular bases of these phenotypes was previously unclear. Our work demonstrates that restricting the genetic inactivation of *Rbfox1* to cINs replicates these findings and identifies that an overall reduction of inhibition in the cortex is the likely cause of these phenotypes. With regard to specific regulatory networks, previous work has shown that Rbfox1 regulates the splicing of ion channels, neurotransmitter receptors and cell adhesion molecules (Fogel et al., 2012; Gehman et al., 2011; Lovci et al., 2013; Voineagu et al., 2011; Weyn-Vanhentenryck et al., 2014; Zhang et al., 2008). However, the extent to which targets of AS vary across cell types and across development has remained unclear. The first hint that transcripts were specifically alternatively spliced in individual neurons was suggested over thirty years ago in *Aplysia* (Buck et al., 1987). Since then few studies have provided examples of particular transcripts being spliced within discrete neuronal populations (Benjamin and Burke, 1994; Fuccillo et al., 2015; Sommer et al., 1990). However, studying how the diversity and regulation of AS patterns is controlled within different cell types has remained technically challenging and to date has mostly been examined in invertebrates (Norris et al., 2014). The striking differences in the Rbfox1 targets found in PV+ versus SST+ cINs provides insight into how a single splicing regulator can differentially modulate gene expression within two related but distinct cell types. Hence, our findings constitute, to our knowledge, the first evidence of distinct cell-type specific AS within closely-related mammalian neuronal types. It will be interesting to investigate the molecular mechanisms underlying these interneuron subtype specific AS events.

Recent work has focused on elucidating the molecular basis for neuronal diversity. In particular, advances in high-throughput genomics and single cell RNA-seq have for the first time provided a fine grain view of the expression of genes in individual neuronal subtypes and across development (Mayer et al., 2018; Mi et al., 2018; Nowakowski et al., 2017; Tasic et al., 2016; 2017; Zeisel et al., 2015). PV+ and SST+ cINs share a common developmental history, have a mutual requirement for key genetic factors including the transcription factors Nkx2-1, Lhx6, Sox6 and Satb1, and arise from a common lineage (Bandler et al., 2017). Our work adds to this by providing a compelling demonstration that beyond transcriptional differences, some of the unique specializations found within specific cINs rely on differential AS networks, that in part are dependent on Rbfox1.

Mutations of *RBFOX1* are known to be associated with ASD, ID and epilepsy (Bhalla et al., 2004; Martin et al., 2007; Sebat et al., 2007). Our understanding of the molecular basis of these disorders has to date relied on global investigations in human brain tissue (Parikshak et al., 2016; Voineagu et al., 2011), constitutive mouse models (Gehman et al., 2011) and knock-down experiments on cultured neurons and cells (Fogel et al., 2012; Lee et al., 2016; Weyn-Vanhentenryck et al., 2014). Our findings, by focusing on the consequences of *Rbfox1* removal in defined cIN classes, illustrate the value of examining the role of disease genes across development and in particular cell types. As such, it will be interesting to further establish the signature of Rbfox1-dependent AS programs within the different neuronal classes that express this gene to gain a clearer image of the causal mechanisms in neurodevelopmental disorders.

## Material and Methods

- Key resources table
- Contact for reagent and resource sharing
- Experimental model and subject details

○ Mouse maintenance and mouse strains
- Method details

○ Immunochemistry and image analysis
○ Confocal imaging and synaptic puncta analysis
○ Seizure susceptibility
○ Electrophysiological recordings
○ Optogenetic stimulation
○ Isolation of cortical interneurons from the developing mouse cerebral cortex
○ Nucleic acid extraction, RNA amplification, cDNA library preparation and RNA sequencing
○ Differential expression and alternative splicing analysis
○ GO analysis
○ Validation of cell-specific Rbfox1-dependent exons by RT-PCR
○ iCLIP analysis
- Quantification and statistical analysis
- Data and software availability

○ Data resources

## CONTACT FOR REAGENT AND RESOURCE SHARING

Please contact GF for reagents and resources generated in this study.

## EXPERIMENTAL MODEL AND SUBJECT DETAILS

### Mouse maintenance and mouse strains

All experimental procedures were conducted in accordance with the National Institutes of Health guidelines and were approved by the Institutional Animal Care and Use Committee of the NYU School of Medicine. Generation and genotyping of *Dlx6a^Cre^* (Yu et al., 2011), *SST^Cre^* (Taniguchi et al., 2011), *Tac1-IRES2-Cre* (referred to as *Tac1^Cre^*) (Harris et al., 2014), *RCE^eGFP^* (Sousa et al., 2009), *Lhx6* BAC transgenic (referred to as *Lhx6::eGFP*) (Gong et al., 2003; Heintz, 2004), *Rbfox1^LoxP/LoxP^* (Gehman et al., 2011), TRE-Bi-SypGFP-TdTomato (Li et al., 2010) and Ai9*^LoxP/LoxP^*, Ai32*^LoxP/LoxP^*, *Rosa-tTa^LoxP/LoxP^*. All mouse strains were maintained on a mixed background (Swiss Webster and C57/ B16). The day of plug is considered as E0.5, the day of birth is considered P0. Information about the mouse strains including genotyping protocols can be found at http://www.jax.org/ and elsewhere (see above references).

## METHOD DETAILS

### Immunochemistry and imaging

Embryos, neonate, juvenile and adult mice were perfused inter cardiac with 4% PFA after being anesthetized either on ice or using Sleepaway IP administration. Brains that were processed for free-floating immunofluorescence were first post-fixed in 4% PFA overnight at 4°C. Brains were sectioned on a Leica vibratome at 50 µm-thickness and stored in a cryoprotecting solution (40% PBS, 30% glycerol and 30% ethylene glycol) at −20°C until use. For immunofluorescence, floating sections were placed in 2 ml tubes for > 30 min at RT in PBS, then blocked for > 1 hr at RT in blocking buffer and incubated for 2-3 days at 4°C with primary antibodies in blocking buffer. Sections were washed 3 × 30 min at RT in PBS, incubated overnight at 4°C with secondary antibodies and DAPI in blocking buffer, washed 3 × 30 min at RT in PBST and once with PBS before being mounted on glass slides.

Brains that were processed for immunofluorescence on slides were post-fixed and cryopreserved following the perfusion and brain harvest. 16µm coronal sections were obtained using Cryostat (Leica Biosystems) and collected on super-frost coated slides, then allowed to dry and stored at −20°C until use. For immunofluorescence, cryosections were thawed and allowed to dry for 5-10 min and rinsed twice in 1x PBS.

They were incubated at room temperature in a blocking solution of PBST (PBS-0.1%Tx-100) and 10% normal donkey serum (NDS) for 60min, followed by incubation with primary antibodies in PBS-T and 1% NDS at 4°C overnight. Samples were then washed 3 times with PBS-T and incubated with fluorescence conjugated secondary Alexa antibodies (Life Technologies) in PBS-T with 1% NDS at room temperature for 60-90min. Slides were then incubated for 30s with DAPI, washed 3 times with PBS-T and once with PBS. Finally, slides were mounted with Fluoromount G (Southern Biotech) and imaged. Primary antibodies are listed in Key Resource Table.

### Confocal imaging and synaptic puncta analysis

Animals were perfused as described above. Post-fixation incubation prior to cryopreservation was skipped. Cryostat sections (16 μm) were subjected to immunohistochemistry as described above. Images were taken within the somatosensory cortex of at least three different sections from three different animals per genotype with a Zeiss LSM 510 and 800 laser scanning confocal microscope. Scans were performed to obtain 4 optical Z sections of 0.33 μm each (totaling ~1.2μm max projection) with a 63x/1.4 Oil DIC objective. The same scanning parameters (i.e. pinhole diameter, laser power/offset, speed/averaging) were used for all images. Maximum projections of 4 consecutive 0.33μm stacks were analyzed with ImageJ (NIH) puncta analyzer plugin (Ippolito and Eroglu, 2010) to count the number of individual puncta consisting of pre-synaptic and post-synaptic markers that are close enough together to be considered a putative synaptic puncta. Synaptic puncta density per image was calculated by normalization to total puncta acquired for each individual channel accounted in each image for each condition. Puncta Analyzer plugin is written by Barry Wark, and is available for download (https://github.com/physion/puncta-analyzer).

### Seizure susceptibility

We investigated the seizure susceptibility according to a modified Racine Scale (Gehman et al., 2011; Racine, 1972). Stage 0, normal behavior; stage 1, immobility; stage 2, mouth and facial movements; stage 3, head bobbing; stage 4, forelimb clonus; stage 5, rearing; stage 6, continuous rearing and falling (tonic-clonic seizures); and stage 7, status epilepticus and/or death. For each animal (aged P12-P14), the score was determined every 5 min for up to 70 mins from time of kainic acid (12mg/kg) intra-peritoneal injection. We used the maximum score of each animal’s behavior at each 5 min interval to determine the average score and standard deviation for both control (*Dlx6a^Cre^;Rbfox1^+/+^;RCE^eGFP^*) (N=5) and *Dlx-cKO* (*Dlx6a^Cre^;Rbfox1^F/F^;RCE^eGFP^*) (N=9).

### Electrophysiological recordings

Slice preparation: Acute brain slices (300 µm thick) were prepared from P17-P60 mice. Mice were deeply anesthetized with euthasol and decapitated. The brain was removed and placed in ice-cold modified artificial cerebrospinal fluid (ACSF) of the following composition (in mM): 87 NaCl, 26 NaHCO_3_, 2.5 KCl, 1.25 NaH_2_PO_4_, 0.5 CaCl, 7 MgCl_2_, 10 glucose, 75 sucrose saturated with 95% O_2_, 5% CO_2_ at pH=7.4. Coronal sections were cut using a vibrating microtome (Leica, VT 1200S). Slices were then incubated at 34-35°C for 30 minutes and then stored at room temperature until use.

Recordings: Slices were transferred to the recording chamber of an up-right microscope (Olympus BX51) equipped with oblique illumination Olympus optics (Olympus) and an infrared camera system (Q-Imaging). Cells were visualized using a 40 or 60X IR water immersion objective. Slices were superfused with preheated ACSF of the following composition (in mM): 124 NaCl, 26 NaHCO_3_, 3 KCl, 1.25 NaH_2_PO_4_, 1.6 CaCl_2_, 2 MgCl_2_, 10 glucose, saturated with 95% O2, 5% CO2 at pH=7.4 and maintained at a constant temperature (31-33 °C). Whole-cell recordings were made from randomly selected tdTomato-positive interneurons or tdTomato-negative pyramidal cells from layer II-III of the somatosensory cortex. Recording pipettes were pulled from borosilicate glass capillaries (Harvard Apparatus) and had a resistance of 3-5 MΩ when filled with the appropriate internal solution, as reported below. Recordings were performed using a Multiclamp 700B amplifier (Molecular Devices). The current clamp signals were filtered at 10 KHz and digitized at 40 kHz using a Digidata 1550A and the Clampex 10 program suite (Molecular Devices). Miniature synaptic currents were filtered at 3 kHz and recorded with a sampling rate of 10 kHz. Voltage-clamp recordings where performed at a holding potential of 0 mV after application of kynurenic acid (3 mM), for current-clamp recordings performed at a holding potential of −65 mV after application of a combination of CNQX (10 μM) and D-AP5 (25 μM). Cells were only accepted for analysis if the initial series resistance was less than 40 MΩ and did not change by more than 20% throughout the recording period. The series resistance was compensated online by at least ~60% in voltage-clamp mode. No correction was made for the junction potential between the pipette and the ACSF.

Passive and active membrane properties were recorded in current clamp mode by applying a series of hyperpolarizing and depolarizing current steps and the analysis was done in Clampfit (Molecular Devices). The highest firing frequency was calculated from a series of 20 pA depolarizing 1 sec long current steps (range 0-520 pA). The cell input resistance was calculated from the peak of the voltage response to a 50 pA hyperpolarizing 1 sec long current step according to Ohm’s law. Analysis of the action potential properties was done on the first spike observed during a series of depolarizing steps. Threshold was defined as the voltage at the point when the slope first exceeds a value of 20 V.s^−1^. Rheobase was defined as the amplitude of the first depolarizing step at which firing was observed. Analysis of miniature inhibitory events was done using Clampfit template search. All values presented in the manuscript are average ± standard error of the mean (SEM) and all the statistical values are obtained doing a standard Student’s t-test, unless otherwise stated (*p ≤ 0.05, **p ≤ 0.01, ***p ≤ 0.005).

Pipette solutions: Solution for voltage-clamp recordings from pyramidal cells (in mM): 130 Cs-methansulfonate, 5 CsCl, 1 HEPES, 0.2 EGTA, 4 MgATP, 0.3 Na-GTP, 5 Phosphocreatine-Tris, 5 QX-314-Cl and 0.3-0.5% biocytin, equilibrated with KOH a pH=7.3. Solution for current clamp recordings from interneurons (in mM): 130 K-Gluconate, 10 KCl, 10 HEPES, 0.2 EGTA, 4 MgATP, 0.3 NaGTP, 5 Phosphocreatine and 0.3-0.5% biocytin, equilibrated with KOH CO2 to a pH=7.3.

### Optogenetic stimulation

Blue-light (470 nm) was transmitted to the slice from an LED placed under the condenser of an up-right microscope (Olympus BX50). IPSCs were elicited by applying single 1 ms blue-light pulses of varying intensities (max. stimulation intensity ~0.33 mW/mm^2^) and directed to layer II-III of the slice in the recording chamber. Light pulses were delivered every 5 seconds increasing the light intensity. The LED output was driven by a TTL output from the Clampex software of the pCLAMP 9.0 program suite (Molecular Devices).

### Isolation of cortical interneurons from the developing mouse cerebral cortex

Cortical interneurons were dissociated from embryonic cortices (E18.5) using the Worthington papain dissociation kit and in accordance with the manufacturer’s guidelines, except for the concentration of papain used during the tissue digestion (9U/mL).

Cortical interneurons were dissociated from postnatal mouse cortices (P4, P8, P12, P22) as described (Hempel et al., 2007). We collected at least 3-7 cKO and 3-5 *wt* brains (see RNA-seq metadata in Supplementary Table 1), and maintained overall balanced numbers of females and males within each condition, in order to avoid sex-related gene expression biases. Following dissociation, cortical neurons in suspension were filtered and GFP+ or TdTomato+ fate-mapped interneurons were sorted by fluorescence activated-cell sorting on either a Beckman Coulter MoFlo (Cytomation), BD FACSAria II SORP or Sony SY3200. Sorted cINs are collected and lyzed in 500µl TRIzol LS Reagent, then thoroughly mixed and stored at −80°c until further totRNA extraction.

### Nucleic acid extraction, RNA amplification, cDNA library preparation and RNA sequencing

Total RNAs from sorted cINs were extracted using TRIzol LS Reagent and PicoPure columns (if less than 20K cells were recovered) or PureLink RNA Mini Kit (if more that 20K cells were recovered), with PureLink DNase for on-column treatment, following the manufacturers’ guidelines. RNA quality and quantity were measured with a Picochip using an Agilent Bioanalyzer. 20ng of total RNA was used for cDNA synthesis and amplification, using NuGEN Ovation RNA-Seq System V2 kit (NuGEN part # 7102). 100 ng of amplified cDNA were used to make a library using the Ovation Ultralow Library System (NuGEN part # 0330). 10 cycles of PCR were run during the amplification step. The samples were pooled and run as 50-nucleotide paired-end read rapid with the Illumina HiSeq 2500 sequencer (v4 chemistry), to generate >50 million reads per sample. Library preparation, quantification, pooling, clustering and sequencing was carried out at the NYULMC Genome Technology Center. qRT-PCR (quantitative RT-PCR) was performed using SYBR select master mix (Thermo-Fisher Scientific) on cDNA synthesized using SuperScript II reverse transcriptase and oligo(dT) primers.

List of RT- and qRT-PCR primers:

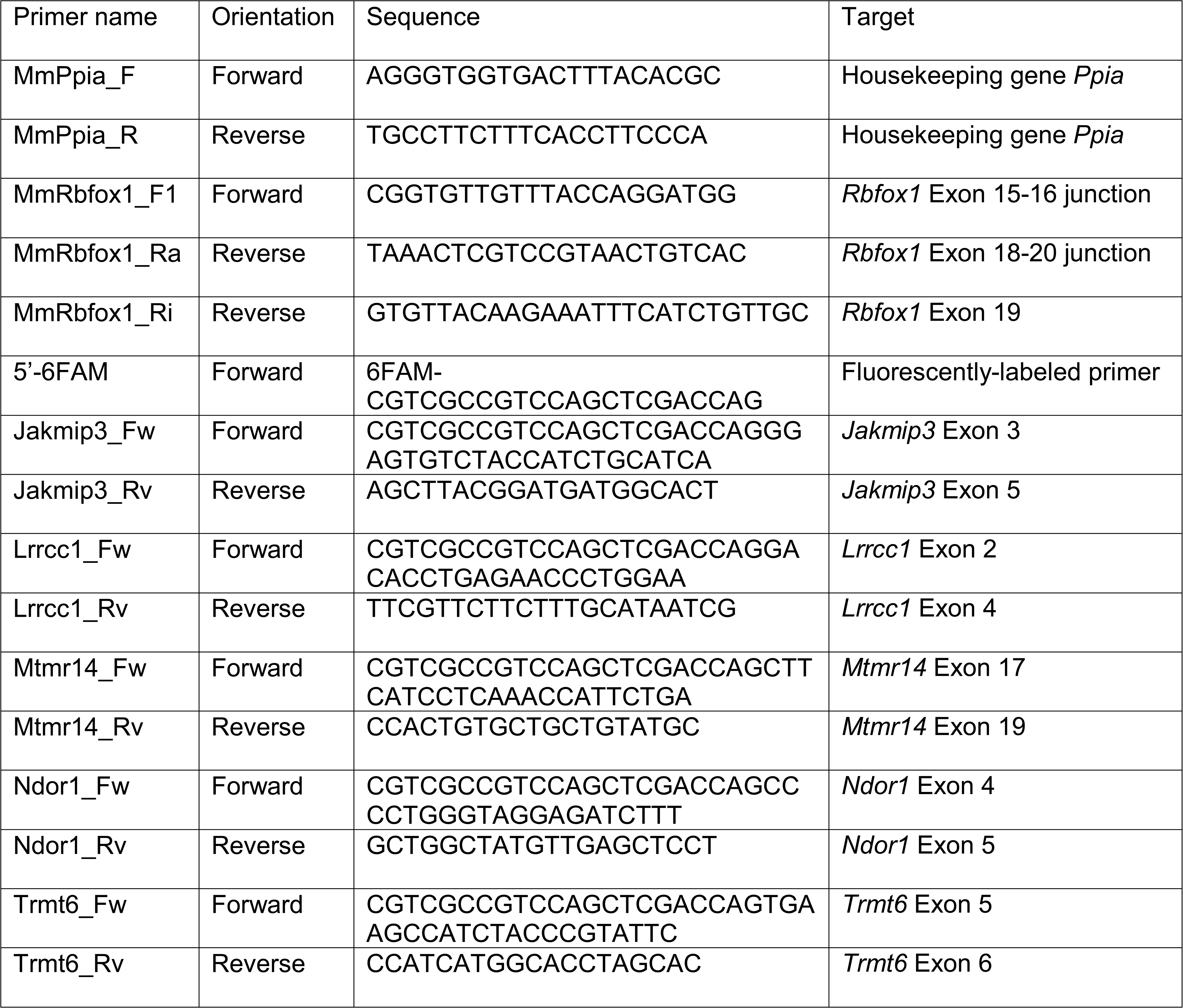

### Differential gene expression and alternative splicing analyses

Downstream sample demultiplexing and computational analysis were performed at the NYULMC Genome Technology Center. All the reads were mapped to the mouse reference genome (GRCm38.74/mm10) using the STAR aligner (v2.5.0c) (Dobin et al., 2013). Alignments were guided by a Gene Transfer File (Ensembl GTF version GRCm38.74) and the mean read insert sizes and their standard deviations were calculated using Picard tools (v.1.126) (http://broadinstitute.github.io/picard/). The Read Per Million (RPM) normalized BigWig files were generated using BEDTools (v2.17.0) (Quinlan and Hall, 2010) and bedGraphToBigWig tool (v4) (see RNA-seq metadata in Supplementary Table 1).

Transcripts were assembled using cufflinks (v2.0) (Trapnell et al., 2013) and their differential expression analysis was performed using Cuffdiff. We used an event-based tools to study different aspects of splicing. We used rMATS (v3.0.9) (Shen et al., 2014) to study the event types such as skipped exons (SE), alternative 3’ splice sites (A3SS), alternative 5’ splice sites (A5SS), mutually exclusive exons (MXE) and retained introns (RI). rMATS uses a counts-based model, detects alternative splicing events using both splice junction and exon body counts and ascribes an exon inclusion level value ψ for each event in each condition. It then determines the differential |∆ψ| value across conditions (cut-offs for significance were placed at *p*-val<0.05 and |∆ψ|≥0.1, unless stated otherwise). To compare the level of similarity among the samples and their replicates, we used two methods: classical multidimensional scaling or principal-component analysis and Euclidean distance-based sample clustering. The downstream statistical analyses and generating plots were performed in R environment (v3.1.1) (http://www.r-project.org/).

### Validation of cell-specific Rbfox1-dependent exons by RT-PCR

Total RNAs from sorted cINs were extracted as described above and at least three independent biological replicates were used in each experiment. RT-PCR validation of regulated exons was performed as described before (Han et al., 2014). After denaturation, samples were run on 10% Novex™ TBE-Urea Gels (ThermoFisher). Gels were directly scanned by ChemiDoc™ Imaging System (Bio-Rad) and quantified by ImageStudio program (Licor).

### GO analysis

We performed GO analysis using the DAVID online Bioinformatics Resources 6.8 (Huang et al., 2008).

### iCLIP analysis

We analyzed the iCLIP data RNA-seq reads from (Damianov et al., 2016) (Rbfox1-HMW-forebrain) available on GEO (GSM1835189). All of the reads were mapped to the mouse reference genome (GRCm38.74/mm10) as above, and duplicate reads were removed using Picard tools. Peak calling was performed using MACS (v1.4.2) (Zhang et al., 2008) and peak count tables were created using BEDTools (Quinlan and Hall, 2010). Differential binding analysis was performed using DESeq2. ChIPseeker (v1.8.0) (Yu et al., 2015) and R package were used for peak annotations and. The RPM normalized BigWig files were generated using MACS and wigToBigWig.

## QUANTIFICATION AND STATISTICAL ANALYSIS

In all figures: *, *p-*value<0.05; **, *p-*value<0.01; ***, *p-*value<0.001; ****, *p-*value<0.0001. Statistical analyses for differential gene expression and alternative splicing changes were performed using DESeq, DEX-seq and rMATS. To quantify the layer distribution and density of various populations of cortical interneuron, the proportion of interneurons of given subtypes over the total number of fate-mapped interneurons across cortical layers was calculated in at least three cryostat tissue sections from a minimum of three individual brains. Thus, percentages presented in Figure 2 and 4 were calculated by dividing the number of markerX+/reporter+ neurons in each layer (eg, layer I, layerII/III, layerIV and layerV/VI) by the total number of reporter+ neurons. Percentages were compared with repeated t-tests in *GraphPad Prism*, and means ± (standard deviation, SD) are represented.

## DATA AND SOFTWARE AVAILABILITY

The accession number for the RNA sequencing data reported in this paper is NCBI GEO: [*TBD*].

## Author contributions

XJ, BW and GF conceived and developed the methodology and project. XJ, BW and NY performed the wet-lab experiments: histology, behavior, imaging, image analysis. AKJ and XJ analyzed the RNA-seq experiments. EF performed and analyzed the target validation experiments. GQ and MJN performed the electrophysiological experiments. BR has provided guidance on the electrophysiological investigations. XJ and BW analyzed and interpreted the results. XJ, BW and GF wrote the paper with review and editing provided by all authors. Project administration: XJ and GF. Funding acquisition: XJ and GF.

## Acknowledgements

We thank Prof. Doug Black (UCLA) for the Rbfox1 mAb and Prof. Südhof (Stanford) for the Syt2 Ab. We thank the cores and shared resources of the NYULMC division of advanced research technologies and their personnel for their technical assistance: Mouse genotyping core (Jiali Deng and Jisen Dai), Cytometry and Cell sorting core (Kamilah Ryan, Keith Kobylarz, Yulia Chupalova and Michael Gregory), Genome Technology Center (Yutong Zhang, Olga Aminova, Adriana Heguy) and Applied Bioinformatics Laboratory (Aristotelis Tsirigos), which is supported in part by grant UL1 TR00038 from the National Center for Advancing Translational Sciences (NCATS), NIH. CCSC and GTC are supported by the Cancer Center Support Grant, P30CA016087, at the Laura and Isaac Perlmutter Cancer Center. We are grateful to Celine Vuong and Doug Black for their comments on the manuscript and to Chia-Ho Lin for her assistance with the iCLIP data analysis. Work in the GF lab is supported by the following NIH grants: R01 NS081297, R01 MH071679, R01 MH111529, P01 NS074972 and by the Simons Foundation (SFARI grant 274578). XHJ was supported by EMBO (ALTF 303-2010), HFSP (LT000078/2011-L), the Bettencourt-Schueller Foundation and by the NIH (K99 MH107684). BW was supported by a training grant (NIH, 5T32NS086750).

